# *Lactobacillus acidophilus* ameliorates inflammatory bone loss under post-menopausal osteoporotic conditions via preventing the pathogenic conversion of gut resident pTregs into Th17 cells

**DOI:** 10.1101/2025.03.20.644481

**Authors:** Asha Bhardwaj, Leena Sapra, Abhay Tiwari, Pradyumna K. Mishra, Rupesh K. Srivastava

**Affiliations:** Translational Immunology, Osteoimmunology & Immunoporosis Lab (TIOIL), An ICMR Collaborating Centre for Excellence on Bone Health, Department of Biotechnology, All India Institute of Medical Sciences (AIIMS), New Delhi-110029, India; Centre for Rural Development & Technology, Indian Institute of Technology (IIT), New Delhi, India; Department of Molecular Biology, ICMR-National Institute for Research in Environmental Health, Bhopal, MP, 462001, India

**Keywords:** pTregs, tTregs, exFoxp3 Tregs, Th17 cells, *Lactobacillus acidophilus*, Butyrate

## Abstract

Research in the past decade has elucidated the explicit role of immune system in the pathophysiology of osteoporosis. Recent studies revealed more details of bone immune interaction and explored various safe and effective immunomodulatory therapies, such as probiotics for the prevention and management of osteoporosis. As a result, various immune factors have continuously been discovered to play specific roles in maintaining bone homeostasis. The role of the Tregs is already well established in the context of post-menopausal osteoporosis (PMO). While Foxp3^+^ Tregs are mostly matured in the thymus (tTregs), some are also produced from Foxp3^−^CD4^+^ T-cell precursors in the peripheral tissues (pTregs). Intriguingly, the specific role of pTregs or tTregs in PMO is still warranted. Foxp3 is required for the suppressive function of Treg cells, but its stability has been debated. Here, we reveal that RORγT^-^FoxP3^+^pTregs (in the intestine and bone marrow) but not FoxP3^+^tTregs show plastic behavior and loss of FoxP3 expression transdifferentiate pTregs into ex-Foxp3 Tregs (i.e., osteoclastogenic Th17 cells), under estrogen-deficient PMO conditions. Interestingly, *Lactobacillus acidophilus* (LA) supplementation in a butyrate-mediated manner stabilizes the expression of FoxP3 in pTregs. Furthermore, butyrate-primed pTregs are more potent in inhibiting osteoclastogenesis than the control pTregs. Altogether, our findings for the first time reveal that pTreg-Th17 cell axis do play a pivotal role in the pathophysiology of PMO, and probiotic-LA mitigates PMO-induced bone loss via augmenting the immunoporotic potential of pTregs.

**Graphical abstract:** RORγT^-^FoxP3^+^pTregs (in the intestine and bone marrow) but not FoxP3^+^tTregs show plastic behavior and loss of FoxP3 expression transdifferentiate pTregs into ex-Foxp3 Tregs (i.e. osteoclastogenic Th17 cells), under estrogen deficient PMO conditions. Interestingly, *Lactobacillus acidophilus* (LA) supplementation in a butyrate-mediated manner stabilizes the expression of FoxP3 in pTregs to attenuate inflammatory bone loss associated with PMO.

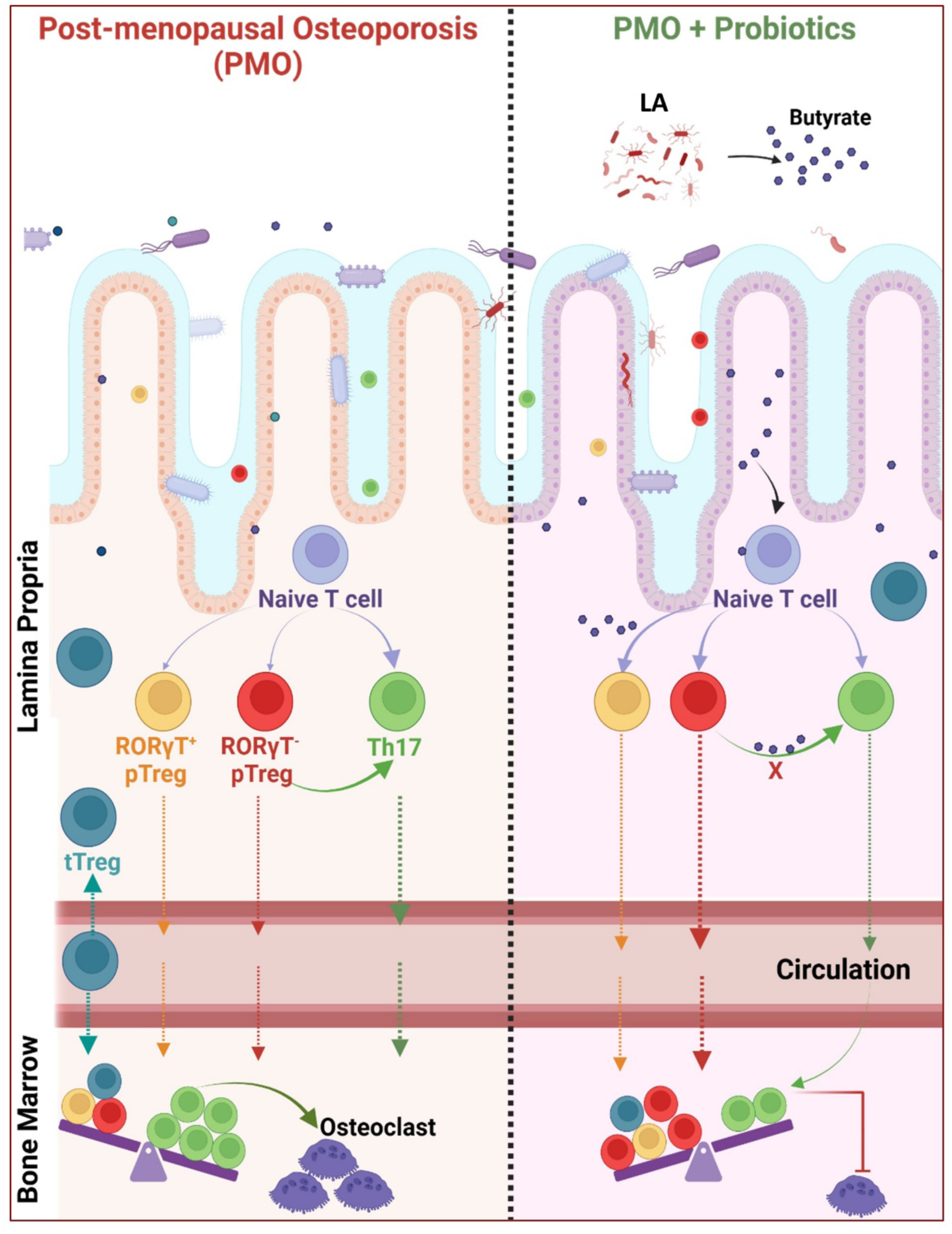

## 1.0 Introduction

Osteoporosis is the most common skeletal bone disease characterized by decreased bone strength and quality, resulting in an increased risk of fractures ^1^. Postmenopausal women are more likely to have osteoporosis than other groups of people worldwide, primarily due to the loss of estrogen after menopause, which promotes bone resorption. Although currently available medications for PMO are effective, patients’ commitment to therapy is hampered by various side effects ^2–5^. Therefore, there is an imperative need for advanced treatments with fewer side effects for the management of post-menopausal osteoporosis (PMO).

Recent studies have evidenced that immune cells are pivotal players in the pathophysiology of osteoporosis. Srivastava et al. (2018, 2021, 2022, 2023) proposed the novel field of “Immunoporosis”, defining and characterizing the increasing role of immune cells in the pathophysiology of osteoporosis^5–8^. The functions of regulatory T cells (Tregs) and Th17 cells in maintaining bone homeostasis have been thoroughly explained in this respect. Tregs are a functionally unique subpopulation of T cells actively maintaining bone homeostasis ^9–13^. Osteoclastogenesis and bone resorption are suppressed by Treg cells in a cytokine-dependent manner through the secretion of anti-inflammatory or immunosuppressive cytokines such as tumor growth factor (TGF)-β and interleukin (IL)-10 ^14–16^. Th17 cells, on the other hand, promote osteoclast differentiation in an IL-17 and RANKL-dependent manner. Targeting Tregs and Th17 cells to regulate pathological immune responses in clinical settings has also been the subject of numerous studies in recent years. Tregs enhancement tactics, which involve increasing the number of antigen-specific or polyclonal Foxp3^+^ Tregs and bolstering their suppressive function, can be useful in managing various inflammatory and autoimmune disorders ^17^. There is evidence from multiple studies indicating that there are two distinct subsets of Treg cells in terms of development: thymus-derived Tregs (tTregs) and peripherally derived Tregs (pTregs), which develop in the thymus as naïve CD4^+^ T cells but become Foxp3 expressing cells in peripheral tissues under the influence of microbiota and their metabolites. A significant portion of pTreg cells in the colon (∼65%) and small intestines (∼35%) also express RAR-related orphan receptor γT (RORγT) and are primarily activated by microbiological cues ^18^.

As there are different subsets of Tregs, we need to better understand which Treg subset can be a therapeutic target in osteoporosis. Since pTregs develop inside the mature T-cell compartment by a variety of stimuli, including probiotics and their metabolites, they represent a viable target population. Nonetheless, it is still unknown to what extent pTregs aid in regulating bone health and whether probiotics and/or their metabolites can attenuate inflammatory bone loss in PMO by regulating the pTreg-Th17 cell axis.

Here, we demonstrate that PMO impairs pTreg’s generation while not affecting the tTregs. The inflammatory environment in PMO stimulates the conversion of RORγT^-^ pTregs to exFoxP3 Th17 cells, thereby decreasing their levels in PMO along with enhancing the population of Th17 cells. We further studied the effect of probiotics in restoring pTregs. We employed the probiotic *Lactobacillus acidophilus* (LA) in the current study as it is one of the most common probiotics with known immunomodulatory and osteoprotective properties ^11^. We observed that LA administration restored the levels of pTreg-1 in the intestine and BM of osteoporotic mice. We further revealed that the levels of short-chain fatty acids (SCFAs) are considerably altered in PMO, and these alterations lead to changes in immune response in the intestine and BM. LA enhances the production of butyrate, which further promotes the development of pTreg-1 by attenuating their conversion to Th17 cells. Furthermore, butyrate-primed pTregs are more potent in inhibiting osteoclastogenesis, thereby preventing bone resorption. To the best of our knowledge, this is the first study that demonstrates LA’s immunomodulatory and osteoprotective potential via its ability to reduce the conversion of RORγT^-^ pTregs to Th17 cells. Our study thus provides novel insights into how dietary interventions such as probiotics can be employed to prevent inflammatory bone loss in PMO via modulating the immunoporotic pTregs.

## 2.0 Material and Methods

### 2.1. Reagents and Antibodies

The following antibodies and kits were purchased from eBioscience (San Diego, CA, USA): PE-Cy7 Anti-Mouse-NRP (25-3041), PerCp-Cy5.5 Anti-Mouse-CD4 (550954), PE Anti-Human/Mouse-Rorγt (12-6988), APC Anti-Mouse/Rat-Foxp3 (17-5773), Foxp3/Transcription factor staining buffer (0-5523-00), Anti-IL-4 (clone: 11B11), Anti-IFN-γ (clone: XMG1.2), and RBC lysis buffer (00-4300-54). The following ELISA kits and reagents were procured from BD (Franklin Lakes, NJ, USA): Mouse TNF-α (OptEIA™-560478) and Mouse IL-6 (OptEIA™-555240). Acid phosphatase leukocyte (TRAP) kit (387A), and retinoic acid (RA) (R2625) were brought from Sigma (St. Louis, MO, USA). Macrophage-colony stimulating factor (M-CSF) (300-25) and receptor activator of nuclear factor κB-ligand (sRANKL) (310-01), murine IL-2 (AF-212-12) and human TGF-β1 (AF-100-21C) were purchased from PeproTech (Rocky Hill, NJ, USA). Roswell Park Memorial Institute (RPMI)-1640 and α-Minimal essential media (MEM) were purchased from Gibco (Grand Island, NY, USA). *Lactobacillus acidophilus* was procured from the American Type Culture Collection (ATCC), USA.

### 2.2 Animals

Female BALB/c mice aged 8-12 weeks were used for *in vitro* and *in vivo* experiments. Mice were housed under specific pathogen-free conditions in the animal facility of the All India Institute of Medical Sciences (AIIMS), New Delhi, India, and given sterilized food and autoclaved drinking water ad libitum. Following anesthesia with xylazine (5-16 mg/kg) and ketamine (100-150 mg/kg), mice were either sham-operated or ovariectomized (ovx). Mice were randomly allocated to three groups: sham, ovx, and ovx + *Lactobacillus acidophilus* (LA). Mice in the ovx + LA group were orally gavaged with 100 µl (10^9^ CFU) of LA suspension daily for 45 days, starting one week post-surgery. At the end of the experiment, mice were euthanized, and several tissues, including blood, bone, mesenteric lymph nodes (MLN), Peyer’s patches (PP), and intestine, were collected. All medical procedures were approved by the Institutional Animal Ethics Committee of AIIMS, New Delhi, India (85/IEAC-1/2018).

### 2.3 Analysis of Short Chain Fatty Acids (SCFAs)

High-performance liquid chromatography (HPLC) was performed for SCFA analysis according to our previous publication ^19^ . Briefly, the first 1 ml of Milli Q and 100 µl of HCl were added to 300 mg of the fecal sample. The sample was completely homogenized with a vortex (2-3 minutes) and placed for 20 minutes. The sample was then centrifuged at 13850 g for 10 minutes (4^0^ C). After centrifugation, the resulting supernatant was transferred to the 2 ml Eppendorf. The supernatant was supplemented with diethyl ether (600 µl) with continued extraction for 20 minutes. After 20 minutes of extraction, the sample was centrifuged at 850 g for 5 minutes (4^0^ C). 400 µl of the supernatant was collected and further supplemented with 500 µl NaOH with continued extraction for 20 minutes. Subsequently the sample was centrifuged at 850 g for 5 minutes. The resulting ether layer was discarded, and the remaining aqueous layer (450 µl) was collected. 300 µl HCl was added to the aqueous layer, and the sample was immediately filtered using the 0.22-micron filter. The resultant sample was run on the HPLC machine. HPLC conditions were: Xterra C18 column (250 mm X 4.6 mm X 3.5 µm); mobile phase, 10 mM H_2_SO_4_ (Isocratic gradient); flow rate, 0.6 ml/min; run time: 11 minutes, wavelength: 210 nm. The SCFA level was determined using the standard calibration curves.

### 2.4 Scanning Electron Microscopy (SEM)

SEM was used to examine the cortical region of femoral bones as per our previous publication ^20^. Briefly, bone samples were maintained in 1% Triton-X-100 for two to three days before being moved to 1% PBS buffer for the duration of the final study. After making the bone slices, the samples were dried under an incandescent lamp before being sputter-coated. Then, using a Leo 435 VP microscope with a 35-mm photographic system, bones were scanned. Digital photos of SEM images taken at a magnification of 100 were used to capture the finest cortical region.

### 2.5 Micro-computed tomography (µ-CT) Measurements

µ-CT scanning and analysis were performed using an *in vivo* X-ray SkyScan 1076 scanner (Aartselaar, Belgium) as per our previous publications ^20^ . Briefly, samples were properly orientated in the sample holder and scanned at 50 kV, 204 mA, with a 0.5-mm aluminum filter. The reconstruction method was carried out using software named NRecon. Following reconstruction, an ROI was formed in the secondary spongiosa 1.5 mm from the distal margin of growth plates using 100 slices. This ROI was then processed by the CTAn program to determine the various micro-architectural properties of bone samples.

### 2.6 Tissue Preparation

Immune cells were isolated from the small intestine (SI) and large intestine (LI) as per our previous publication ^19^. Briefly, after harvesting, MLN, SI, and LI were collected. MLN was minced using frosting end slides, and the resulting suspension was passed through the 70 µM cell strainer. Cells were then processed for flow cytometry. To isolate immune cells from SI, first, PPs were removed and processed similarly to the MLN. After the excision of PP, SI was cleaned and cut into small pieces. Tissue pieces were shaken at 220 rpm for 20 minutes at 37 °C in Hank’s balanced salt solution (HBSS) containing 1mM dithioerythritol for the removal of intraepithelial lymphocytes (IELs). The resulting IELs were processed for cell staining. The remaining tissue was further processed at 37 °C in HBSS containing 1.3 mM ethylenediaminetetraacetic acid (EDTA) with shaking at 220 rpm for 30 minutes to remove epithelial cells. To isolate cells from the lamina propria (LP), the remaining tissue was washed and digested with RPMI containing collagenase type 1 (260 U/ml) by shaking at 220 rpm for 45 min at 37 °C. After tissue digestion, lymphocytes were harvested using a 44% and 67% Percoll gradient purification method. To isolate lymphocytes from the LI, the caecum was removed, and the colon was processed similarly to the SI.

### 2.7 Flow cytometry

Cells were harvested from bone marrow (BM), MLN, PP, and intestine (SI and LI) and stained for Treg and Th17 cells. Cells were initially surface-stained for anti-CD4-PerCPcy5.5 and anti-NRP1-PECy7 with incubation for 30 minutes in the dark. Cells were then fixed and permeabilized for intracellular staining with the help of a 1X-fixation-permeabilization buffer in the dark for 30 minutes. After permeabilization, cells were stained for anti-FoxP3-APC and anti-RORγT-PE for 45 minutes. Cells were acquired on BD LSR fortessa (USA), and data was analyzed with the help of Flowjo-10 software (BD, USA). The gating strategy was adopted as per the experimental requirements.

### 2.8 *In-vitro* differentiation of gut resident Tregs (GTregs)

Naïve CD4^+^ T cells were negatively selected using the T cell enrichment cocktail (BD) from the MLN of 8 to 12-week-old mice and seeded in the plate coated with anti-CD3 (10 µg /ml) and anti-CD28 (2 µg/ml). Cells were further stimulated with TGF-β1 (5 ng/ml), IL-2 (10 ng/ml) anti-IL-4 (5 µg/ml), and anti-IFNγ (5 µg/ml) in the +/-of butyrate (0.3 mM) and IL-6 (30 ng/ml) for 5 days. On day 5, cells were harvested, and flow cytometry was performed to analyze pTregs (CD4^+^Foxp3^+^NRP^-^) and tTregs (CD4^+^Foxp3^+^NRP^+^). For the generation of RORγT^+^ pTregs, 1 nm retinoic acid was added in the above-mentioned conditions, and flow cytometry was performed to analyze the CD4^+^Foxp3^+^RORγT^+^ cells.

### 2.9 Coculture of gut resident pTregs with Bone Marrow Cells (BMCs) for Osteoclastogenesis

Gut resident pTregs were cocultured with osteoclasts according to our previous publication^19^. BMCs were obtained by flushing the femoral bones of mice with complete α-MEM media (10% heat-inactivated FBS) and cultured overnight in a T-25 flask in complete α-MEM media consisting of MCSF at the 35-ng/ml concentration following RBC lysis with 1X RBC lysis buffer. The next day non-adherent cells were collected and co-cultured 50,000 per well with pTregs or butyrate-primed pTregs in different cell ratios (BMC: pTregs; 1:10, 1:5, and 1:1) in 96 well plate in α-MEM media supplemented with M-CSF (30 ng/ml) and RANKL (100 ng/ml) for 4 days. 50% of the media was replenished on the 3^rd^ day. At the end of the experiment, TRAP staining was performed as per the manufacturer’s instructions. Briefly, the cells were washed twice with IX PBS and fixed with the fixative solution consisting of citrate, acetone, and 3.7% formaldehyde solution for 10 minutes. After fixation, cells were washed twice with 1X PBS and then stained with TRAP staining solution with incubation for 5-15 minutes in the dark at 37°C. An inverted microscope (Eclipse, TS100, Nikon, and EVOS, Thermo Scientific, Waltham, MA, USA) was used to count and image multinucleated TRAP-positive cells with more than 3 nuclei (classified as osteoclasts). Osteoclast area and number is estimated with ImageJ software (NIH, USA).

### 2.10 Coculture of gut resident pTregs with BMCs for Osteoblastogenesis

Gut resident pTregs were cocultured with osteoblasts according to our previous publication^19^. Briefly, BMCs were obtained by flushing the femoral bones of mice with complete α-MEM media and seeded 50,000 per well in the presence of osteogenic induction media (OIM) consisting of α-MEM media with ascorbic acid (50 µg/ml) and β-glycerophosphate (10 mM) for 24 hours. On the 2^nd^ and 3^rd^ day, 50% and 80% of the media were replaced with fresh OIM, respectively. The following day, complete media was replaced with fresh OIM and BMCs were cocultured with the pTregs or butyrate-primed pTregs in different cell ratios (BMC: pTregs; 1:10, 1:5, and 1:1). On day 7 alkaline phosphatase (ALP) staining was executed to analyze osteoblastogenesis with media replenishment on every 3^rd^ day. For ALP staining, cells were fixed with 10% formaldehyde for 10 minutes. After fixation, cells were washed twice with 1X PBS. 50 µl nitro blue tetrazolium (NBT)/ 5-bromo-4-chloro-3-indolyl-phosphate (BCIP) substrate was added to each well and the plate was incubated at 37^0^ C until the stain developed.

### 2.11 Statistical Analysis

Statistical analyses have been performed with the help of GraphPad Prism version 8.0 (San Diego, California, USA). All of the data values are expressed as mean ± SEM. Statistical differences between the two groups were evaluated using an unpaired parametric Student’s t-test. Statistical significance for the specified groups was defined as p≤ 0.05 (*p<0.05, **p<0.01, ***p<0.001).

## 3.0 RESULTS

### 3.1 LA administration enhances Bone health by modulating GTregs

To study the probable role of LA in modulating the GTregs, we first examined the potential of LA in mitigating bone loss in PMO. To accomplish the same, mice were divided into three groups: sham (control), ovx (ovariectomized), and ovx + LA. Ovx + LA group was administered daily with 10^9^ CFU of LA for 45 days **(Fig. 1A)**. On day 45, mice were sacrificed, and bones were collected for scanning electron microscopy (SEM) and micro-CT (μ-CT). SEM analysis of the femoral cortical bone sections revealed a large number of resorption pits or lacunae in the ovx group. However, administration of LA to the ovx mice substantially reduced the number of bone resorption pits in comparison to the ovx group **(Fig. 1B)**. µ-CT results revealed that the ovx group has significantly deteriorated micro-architecture of the LV-5 bone as compared to the sham and ovx + LA groups **(Fig. 1C)**. Quantitative examination of trabecular and cortical bones, including key histomorphometric indices generated from 3D bone micro-architecture images is shown in **Table 1** and also has been shown in detail in our previous publication ^11^.

**Figure 1.**
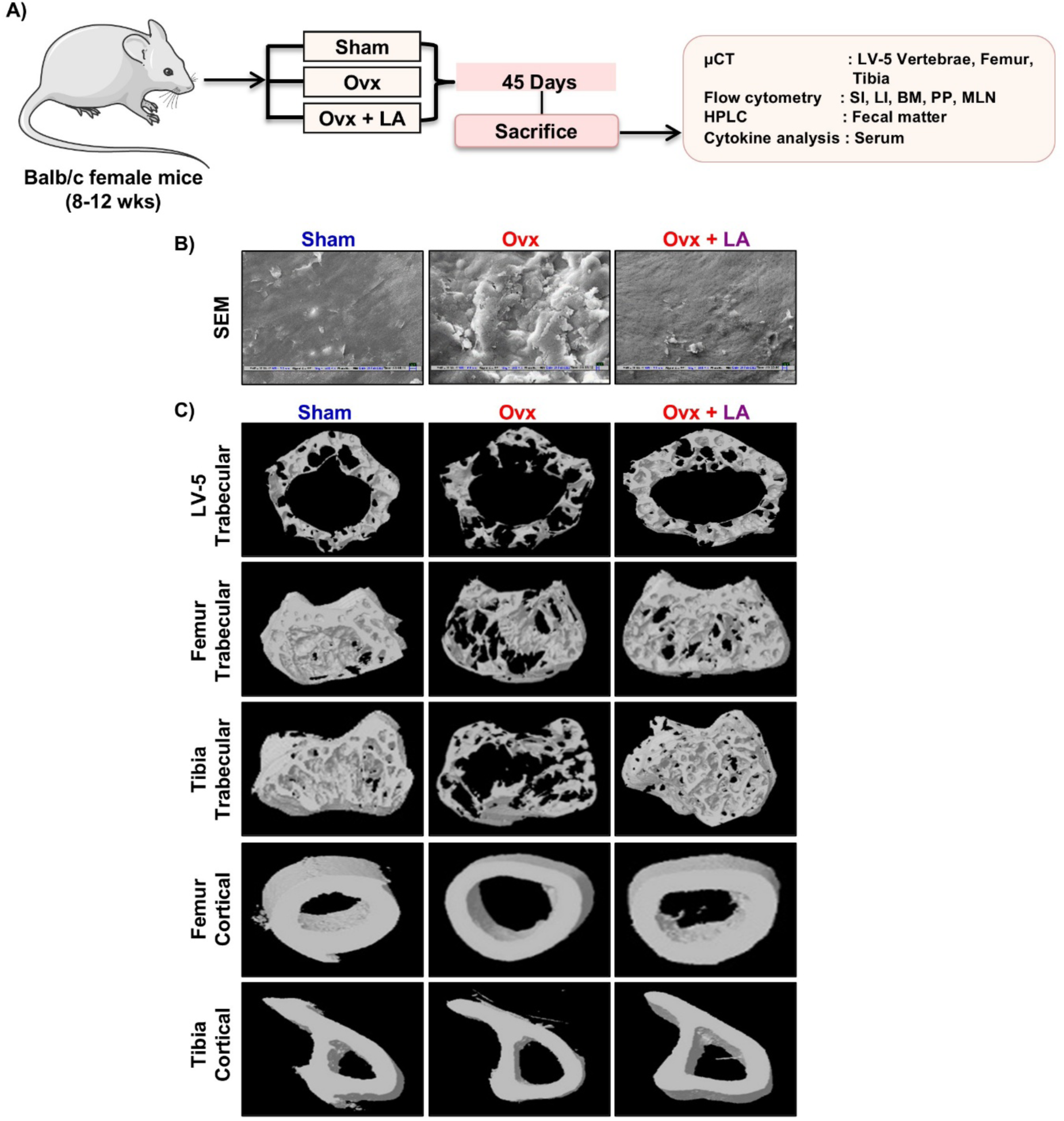
LA administration attenuates bone loss. **A)** *in vivo* experimental work plan. Mice were divided into 3 groups, sham, ovx, and ovx + *Lactobacillus acidophilus* (LA) group which received LA at 10^9^ CFU/day orally reconstituted in drinking water. At the end of 45 days, mice were sacrificed. **B)** 2D SEM images of femur cortical bone. **C)** 3-D μCT reconstruction and histomorphometric indices of LV-5 trabecular, femur trabecular, tibia trabecular, femur cortical, and tibia cortical of all groups.

**Table 1.** Bone histomorphometric parameters in sham, ovx, and ovx + LA group. BV/TV, Bone volume/tissue volume ratio; Tb. Th, trabecular thickness; Tb. No., trabecular number; Conn. Den, connectivity density; Tb. Sep., trabecular separation; Tb. Pf., trabecular pattern factor; Tt. Ar., total cross-sectional area; T. Pm, total cross-sectional perimeter; B. Pm, bone perimeter; Ct. Po, cortical porosity. The results were evaluated by ANOVA with subsequent comparisons by Student t-test for paired or nonpaired data. Values are reported as mean± SEM. Similar results were obtained in two different independent experiments with n=6. Statistical significance was considered as p≤0.05(*p≤0.05, **p≤0.01, ***p≤0.001) with respect to indicated Sham group. Histomorphometric indices of LV-5 trabecular, femur trabecular, tibia trabecular, femur cortical, and tibia cortical of the sham and ovx mice groups.

| Bone Parameters | Sham | Ovx | Ovx + LA |
| --- | --- | --- | --- |
| **LV5** |  |  |  |
| BV/TV (%) | 22.34 ± 0.1773 | 18.52 ± 2.736 | 24.56 ± 0.4118* |
| Tb. Th (mm) | 0.06228 ± 0.001195 | 0.05702 ± 0.001163* | 0.06098 ± 0.001372* |
| Tb. No (mm<sup>-1</sup>) | 3.589 ± 0.06415 | 3.314 ± 0.4238 | 3.783 ± 0.1780 |
| Conn. Den (mm<sup>-3</sup>) | 171.6 ± 31.16 | 149.2 ± 33.65 | 201.7 ± 14.38 |
| Tb. Sp. (mm) | 0.2182 ± 0.004965 | 0.2263 ± 0.01646 | 0.1889 ± 0.007242 |
| Tb. Pf (mm<sup>-1</sup>) | 11.84 ± 0.4165 | 14.43 ± 1.588 | 10.95 ± 0.7084* |
| **Femur Trabecular** |  |  |  |
| BV/TV (%) | 34.94 ± 1.867 | 18.38 ± 1.595** | 3.42 ± 2.387 |
| Tb. Th (mm) | 0.07011 ± 0.002427 | 0.05978 ± 0.001009** | 0.06348 ± 0.001325 * |
| Tb. No (mm<sup>-1</sup>) | 4.978 ± 0.1158 | 3.093 ± 0.2984** | 3.869 ± 0.4341 |
| Conn. Den (mm<sup>-3</sup>) | 361.1 ± 20.23 | 208.7 ± 30.46** | 263.4 ± 39.73 |
| Tb. Sp. (mm) | "0.1560 ± 0.006749 | 0.2668 ± 0.02598** | 0.2066 ± 0.01714* |
| Tb. Pf (mm<sup>-1</sup>) | 2.941 ± 1.837 | 12.05 ± 1.033** | 9.013 ± 1.199 |
| **Tibia Trabecular** |  |  |  |
| BV/TV (%) | 28.34 ± 4.205 | 23.21 ± 2.851* | 20.65 ± 2.559 |
| Tb. Th (mm) | "0.06507 ± 0.0008553 | "0.06251 ± 0.0004700* | "0.06204 ± 0.002084 |
| Tb. No (mm<sup>-1</sup>) | 4.698 ± 0.8038 | 3.577 ± 0.6280 | 3.368 ± 0.5160 |
| Conn. Den (mm<sup>-3</sup>) | 283.0 ± 67.78 | 202.3 ± 41.49 | 184.4 ± 58.20 |
| Tb. Sp. (mm) | 0.2327 ± 0.06970 | 0.2319 ± 0.04062 | 0.2501 ± 0.03684 |
| Tb. Pf (mm<sup>-1</sup>) | 4.363 ± 3.206 | 12.07 ± 1.290* | 13.65 ± 1.275 |
| **Femur Cortical** |  |  |  |
| Tt. Ar (mm<sup>2</sup>) | 1.750 ± 0.02118 | 1.543 ± 0.05400** | 1.683 ± 0.03002* |
| T. Pm (mm) | 5.294 ± 0.1718 | 4.774 ± 0.09391* | 4.978 ± 0.05191* |
| B. Pm (mm) | 8.694 ± 0.2432 | 7.983 ± 0.1248* | 8.676 ± 0.1440** |
| Ct. Po (mm) | 30.64 ± 2.166 | 37.64 ± 1.122* | 32.86 ± 1.368* |
| **Tibia Cortical** |  |  |  |
| Tt. Ar (mm<sup>2</sup>) | 1.399 ± 0.08882 | 1.263 ± 0.06545 | 1.417 ± 0.04056* |
| T. Pm (mm) | 6.132 ± 0.1094 | 5.723 ± 0.1316* | 5.769 ± 0.1410 |
| B. Pm (mm) | 9.477 ± 0.2509 | 8.581 ± 0.1942* | 8.769 ± 0.1243 |
| Ct. Po (mm) | 22.05 ± 3.782 | 26.81 ± 1.304 | 29.66 ± 0.5644* |

Next, to evaluate the role of LA in ameliorating bone loss via its effect on GTregs in PMO, we harvested immune cells from the lamina propria of both small (LP-SI) and large intestine (LP-LI, the prime site of Tregs), intraepithelial lymphocytes from the SI (IEL-SI) and LI (IEL-LI), along with PP, MLN and BM from all the groups and analyzed them for Tregs (CD4^+^Foxp3^+^) (**Supplementary Fig. 1 & 2**). It was observed that the Tregs population was significantly reduced in the LP-SI (p<0.01), LP-LI (p<0.001), IEL-LI (p<0.05), PP (p<0.05), and BM (p<0.01) of ovx mice as compared to the sham **(Fig. 2A-B)**. Remarkably, LA administration significantly enhanced the frequency of Tregs in LP-SI (p<0.01), LP-LI (p<0.01), IEL-SI (p<0.01), IEL-LI (p<0.05) PP (p<0.05), and BM (p<0.01) **(Fig. 2A-B)**.

**Figure 2.**
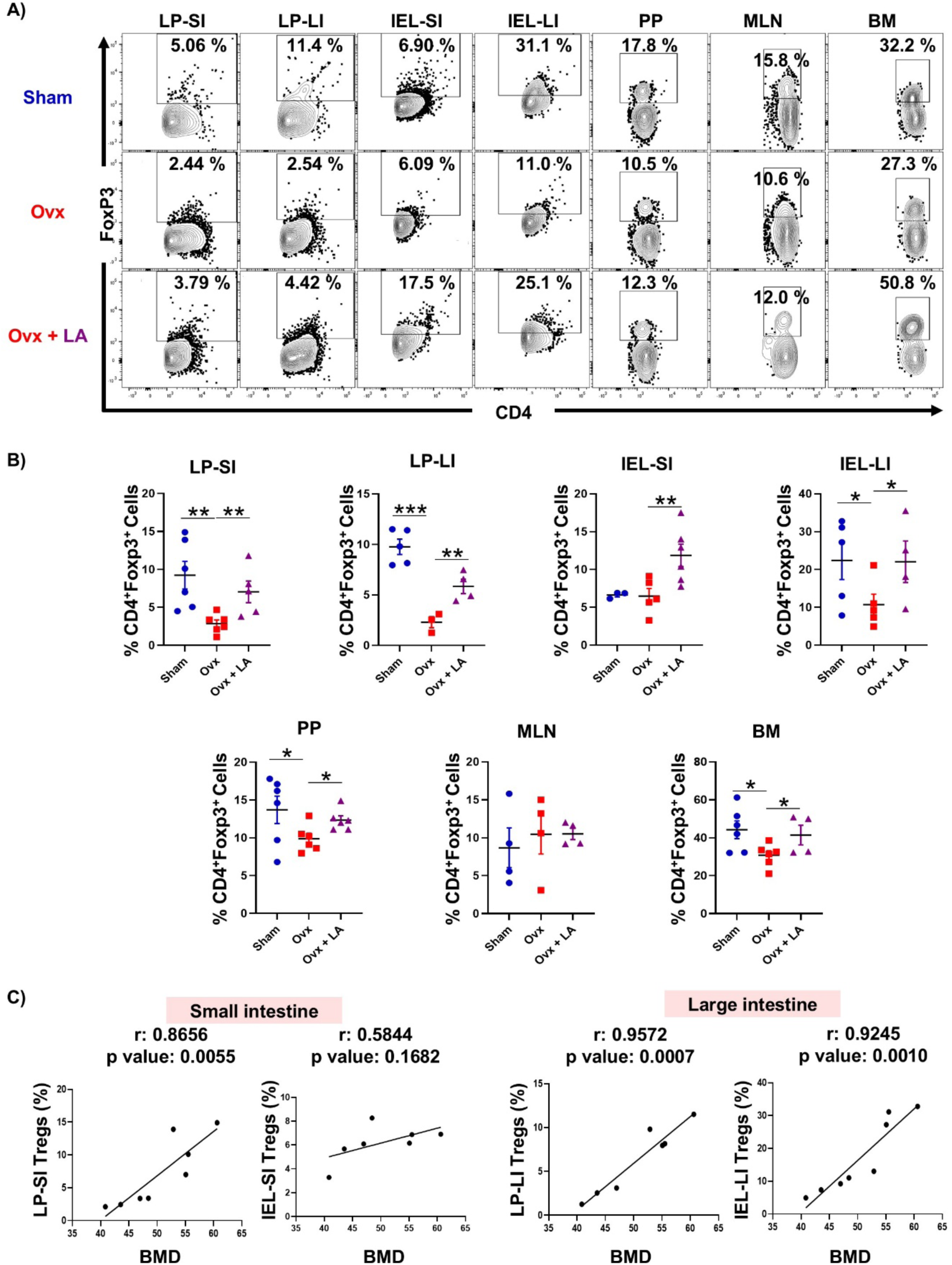
LA administration enhances GTregs *in vivo*. **A).** Cells from lamina propria of the small intestine (LP-SI) and large intestine (LP-LI), intestinal epithelial lymphocytes from the small intestine (IEL-SI) and large intestine (IEL-LI), Peyer’s patches (PP), mesenteric lymph nodes (MLN) and bone marrow (BM) of mice from sham, ovx, and ovx + LA groups were harvested and analyzed by flow cytometry for a percentage of CD4^+^Foxp3^+^Tregs. **B)** Bar graph representing percentages of Tregs in sham, ovx, and ovx + LA. **C)** Correlation graphs depicting the correlation of Tregs from the SI (LP and IEL) and LI (LP-IEL) with the bone mineral density (BMD) of femur trabecular bone. Data are reported as mean ± SEM (n=6). The results were evaluated using the unpaired student t-test. Values are expressed as mean ± SEM (n=6) and similar results were obtained in two independent experiments. Statistical significance was defined as *p ≤ 0.05, **p < 0.01 ***p ≤ 0.001 concerning the indicated mice group.

Furthermore, we observed that the percentage of Tregs in the SI (LP) and LI (LP and IEL) correlated positively with the BMDs of the femur trabecular bones **(Fig. 2C)**. Collectively, our results demonstrate that LA ameliorates inflammatory bone loss in PMO via enhancing GTregs.

### 3.2 LA administration restores the homeostasis of gut resident pTregs and tTregs in ovx mice

Next, to study the effect of LA on two distinct populations of Tregs, the frequency of tTregs and pTregs was analyzed within the Tregs population in various gut tissues (LP-SI, LP-LI, IEL-SI, IEL-LI, PP, and MLN) along with the BM in all the groups. Interestingly it was observed that ovx mice have a significantly decreased percentage of pTregs (CD4^+^Foxp3^+^NRP-1^-^) and an increased percentage of tTregs (CD4^+^Foxp3^+^NRP-1^+^) than sham mice in the gut tissues i.e., LP-SI (p<0.05), LP-LI (p<0.01) and MLN (p<0.01) **(Supplementary Fig. 3 & Fig. 3A-B)**. Surprisingly LA supplementation to ovx mice significantly enhanced the frequency of pTregs along with decreasing the tTregs in the LP-SI (p<0.05), LP-LI (p<0.05), LI-IEL (p<0.05) and MLN (p<0.05). Strikingly LA treatment also enhanced the pTregs and reduced the tTregs in the BM (p<0.05) of ovx mice **(Supplementary Fig. 3 & Fig. 3A-B)**. Additionally, the proportion of pTregs in the LP-SI and LP-LI was positively correlated with the BMDs of femur trabecular bone, while the frequencies of tTregs showed a negative correlation **(Fig. 3C)**. Altogether our results showed that LA administration restores the homeostasis of gut resident pTreg and tTreg cells.

**Figure 3.**
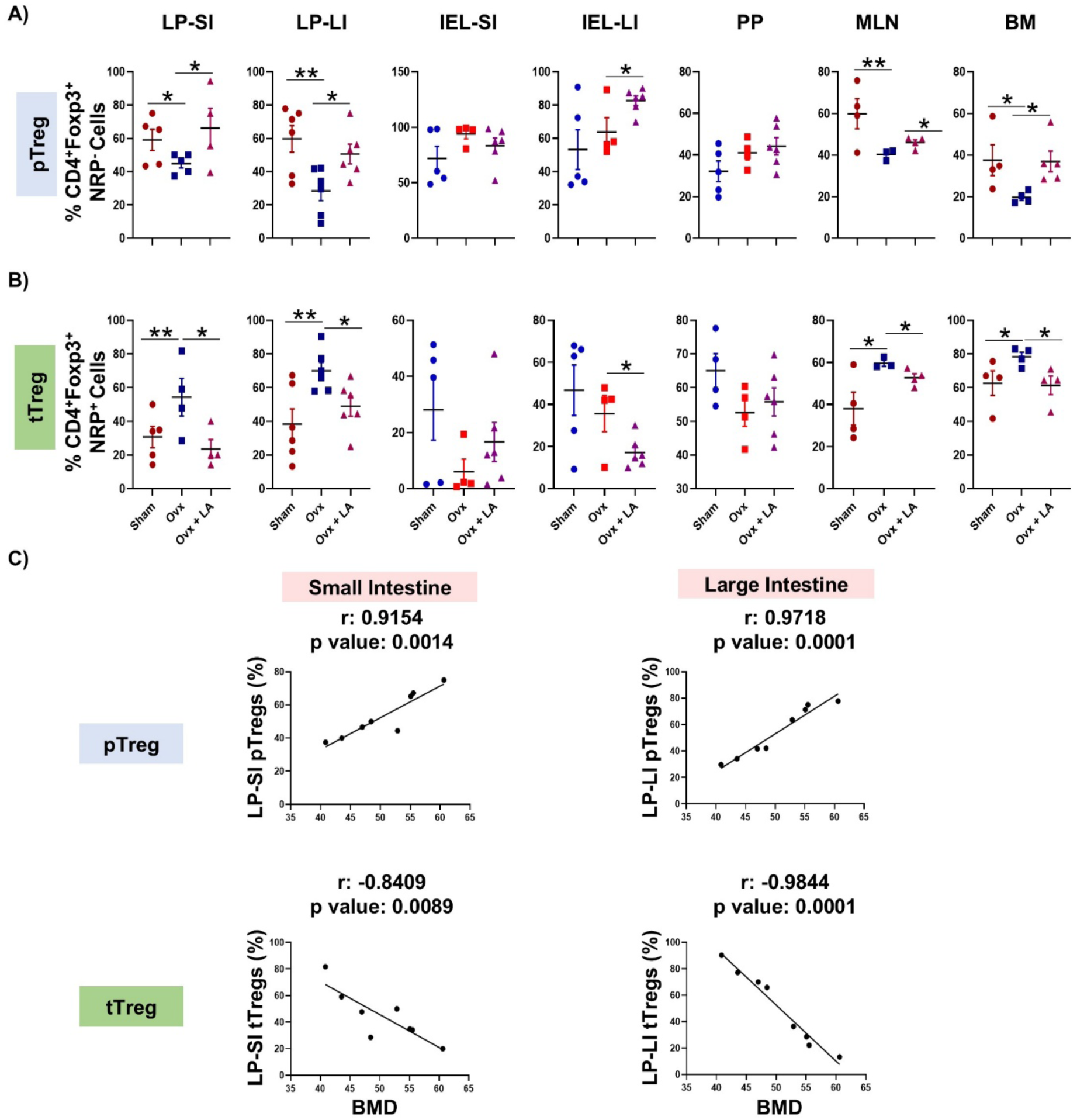
LA administration modulates gut resident pTregs and tTregs *in vivo*. Cells from lamina propria of the small intestine (LP-SI) and large intestine (LP-LI), intestinal epithelial lymphocytes from the small intestine (IEL-SI) and large intestine (IEL-LI), Peyer’s patches (PP), mesenteric lymph nodes (MLN), and BM of mice from sham, ovx and ovx + LA groups were harvested and analyzed by flow cytometry for the percentage of pTregs (CD4^+^NRP^-^Foxp3^+^ cells) and tTregs (CD4^+^NRP^+^Foxp3^+^ cells). **A)** Bar graph representing percentages of pTregs in sham, ovx, and ovx + LA groups. **B)** Bar graph representing percentages of tTregs in sham, ovx, and ovx + LA groups. Data are reported as mean ± SEM (n=6). **C)** Correlation graphs depicting the correlation of pTregs and tTregs from the LP-SI and LP-LI with the bone mineral density (BMD) of femur trabecular bone. The results were evaluated using the unpaired student t-test. Values are expressed as mean ± SEM (n=6) and similar results were obtained in two independent experiments. Statistical significance was defined as *p ≤ 0.05, **p < 0.01 ***p ≤ 0.001 concerning the indicated mice group.

### 3.3 LA administration enhances the differentiation of pTregs in a butyrate-mediated manner

LA enhances the production of short-chain fatty acids (SCFAs)^21^. Thus, to investigate the mechanism of LA-mediated modulation of pTreg and tTreg cells, we next performed HPLC for targeted analysis of the SCFAs (acetate, propionate, and butyrate) in the fecal samples from all the groups of mice. HPLC data demonstrated that ovx mice have significantly decreased amounts of acetate (p<0.05), propionate (p<0.05), and butyrate (p<0.05) in the fecal samples as compared to the controls **(Fig. 4A).** On the other hand, administration with the probiotic LA significantly enhanced the level of the butyrate (p<0.05) in comparison to the ovx mice **(Fig. 4A)**. No significant enhancement was observed in the levels of acetate, and propionate indicating that LA particularly enhanced the levels of butyrate in the ovx + LA group **(Fig. 4A).**

**Figure 4.**
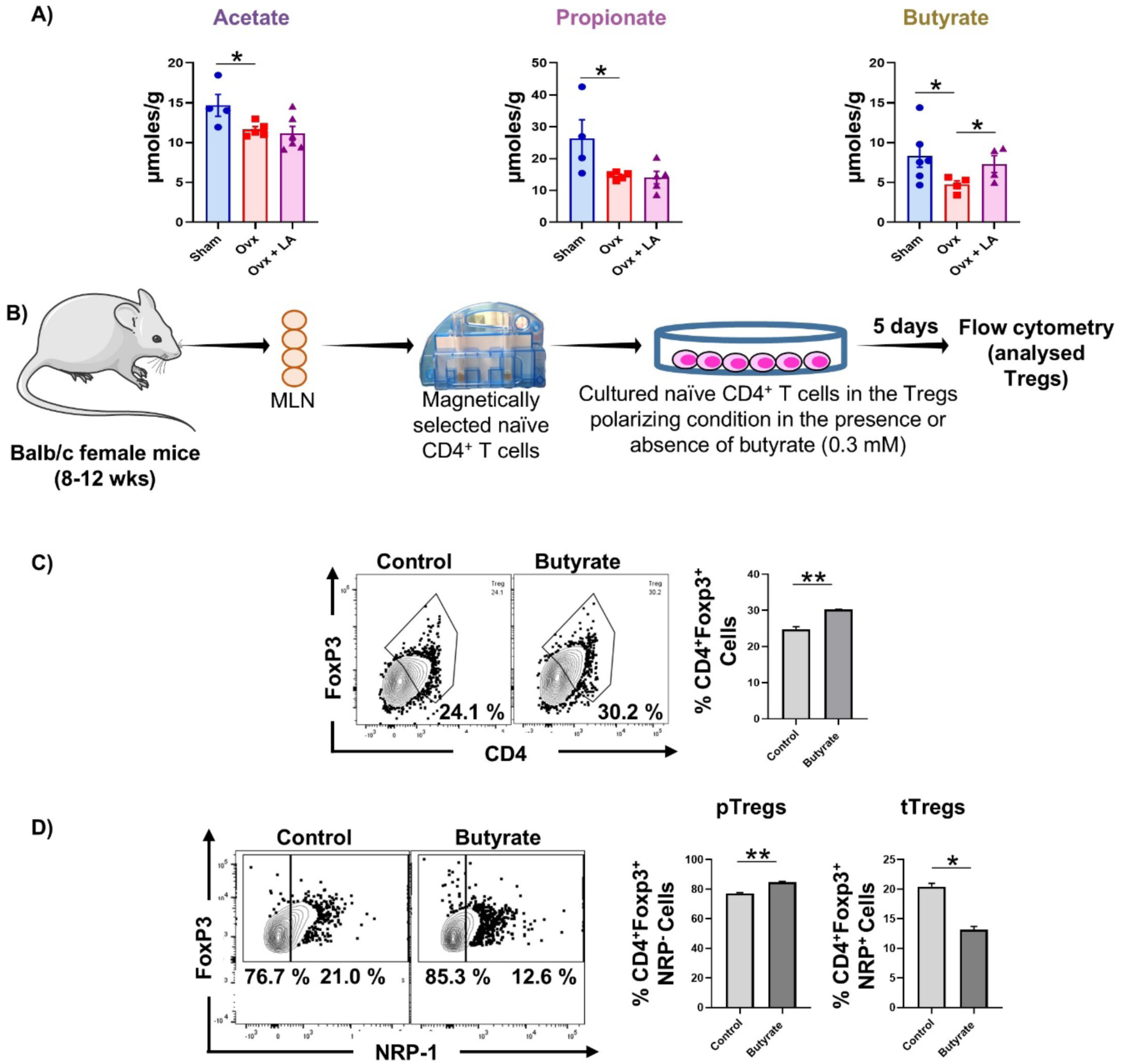
LA administration enhances the level of butyrate in ovx mice. **A)** Short-chain fatty acids (Acetate, Propionate, and Butyrate) were analyzed in the fecal content of all the groups with the help of HPLC. Values are expressed as mean ± SEM (n=6). Statistical significance was defined as p≤0.05, *p≤0.05, **p < 0.01 ***p≤0.001 for the indicated mice group. **B)** Naïve T cells were isolated from mice’s mesenteric lymph nodes (MLN) and induced into Tregs in the presence and absence of butyrate. **C)** Tregs and **D)** pTregs and tTregs were analyzed by flow cytometry in the total induced T cells in the presence and absence of butyrate. The results were evaluated using the unpaired student t-test. Similar results were obtained in two independent experiments. Statistical significance was defined as *p ≤ 0.05, **p < 0.01 ***p ≤ 0.001 concerning the indicated mice group.

Moving ahead we next delineated the effect of SCFAs on the differentiation of the gut resident pTregs *ex vivo*. To accomplish the same, naïve T cells from the MLN (gut-associated lymphoid tissue) were isolated and cultured under Tregs polarizing conditions in the presence or absence of butyrate (0.3M) **(Fig. 4B).** After 5 days of the culture, Tregs were analyzed, and it was observed that butyrate significantly enhanced the frequency of Tregs (p<0.01) as compared to the control **(Fig. 4C).** Interestingly within the Tregs, butyrate treatment significantly increased the differentiation of pTregs (p<0.01) along with inhibiting the differentiation of the tTregs (p<0.05) in comparison to the controls **(Fig. 4D).**

### 3.4 Butyrate inhibits the conversion of pTregs into Th17 cells

Previous studies have shown that under inflammatory conditions (increased levels of IL-6 and decreased levels of IL-10), pTregs can transdifferentiate into ex-FoxP3 Tregs (i.e., Th17 cells) ^22–24^. Thus, we next analyzed the levels of inflammatory cytokines and observed increased levels of inflammatory cytokines IL-6 (p<0.001), TNF-α (p<0.001), and IL-17(p<0.001) along with significantly decreased levels of anti-inflammatory cytokine IL-10 (p<0.001) in the serum of ovx group as compared to the control group **(Figure 5A).** Interestingly, ovx + LA group had significantly decreased levels of inflammatory cytokines, IL-6 (p<0.001), TNF-α (p<0.001), and IL-17 (p<0.001) along with significantly increasing the level of anti-inflammatory IL-10 (p<0.001) in contrast to the ovx group **(Fig. 5A).** These results highlight that the decreased frequency of pTregs in ovx mice might be due to their conversion into exFoxp3 Tregs.

**Figure 5.**
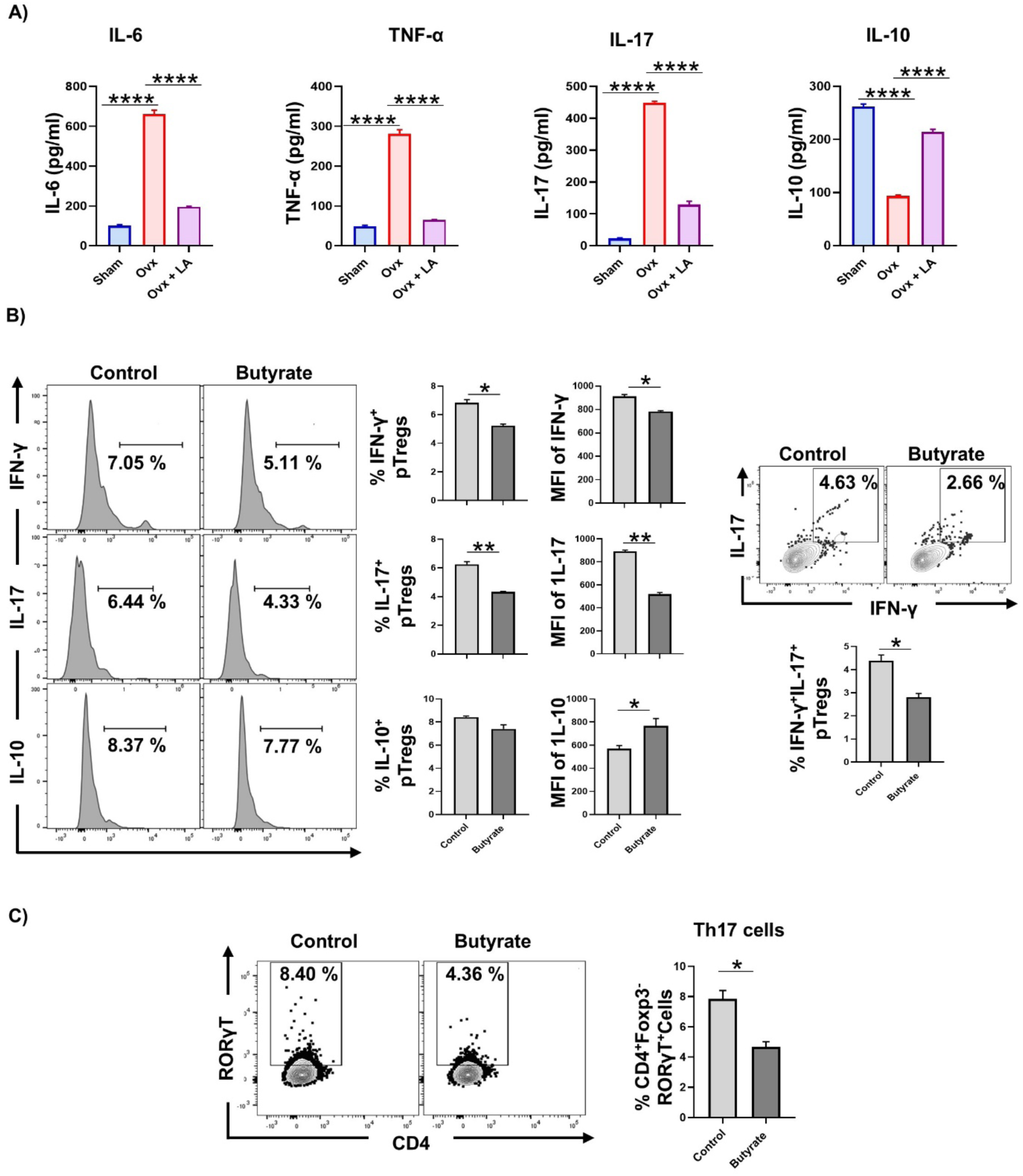
LA administration prevents the conversion of pTregs to Th17 cells. **A)** Inflammatory cytokines were analyzed in serum samples of mice by ELISA. **B)** Analysis of IFN-γ, IL-17, and IL-10 levels in the pTregs population by flow cytometry in the presence and absence of butyrate. **C**) Analysis of Th17 cells in the pTregs populations by flow cytometry in the presence and absence of butyrate. The results were evaluated using the unpaired student t-test. Similar results were obtained in two independent experiments. Statistical significance was defined as *p ≤ 0.05, **p < 0.01 ***p ≤ 0.001 concerning the indicated mice group.

We were next interested in elucidating whether LA in a butyrate-mediated manner can prevent the conversion of pTregs to exFoxp3 Tregs. To accomplish the same, naïve T cells isolated from the MLN were cultured under Tregs polarizing conditions in the presence/absence of butyrate. After 5 days, expression of IFN-γ and IL-17 (markers for exFox3 Tregs) was analyzed via flow cytometry **(Fig. 5B).** We observed that butyrate treatment significantly reduced the production of IFN-γ (p<0.05) and IL-17 (p<0.01) specifically from pTregs, but not from tTregs **(Supplementary Fig. 4)**. Butyrate treatment was further observed to significantly decrease the frequency of IFNγ^+^IL-17^+^ Tregs compared to the control, thereby indicating that butyrate prevents the conversion of pTregs to exFoxp3 Tregs **(Fig. 5B).** Since ex-FoxP3 Tregs have a similar profile as Th17 cells, thus we further analyzed Th17 cells in our *in vitro*-induced Tregs. Interestingly, we observed that butyrate significantly decreased Th17 cells compared to the control, thereby indicating that butyrate prevents the transdifferentiation of pTregs into Th17 cells **(Fig. 5C).**

Moving ahead, we next corroborated our above *ex vivo* findings under *in vivo* pre-clinical settings in ovx mice. For the same, we analyzed Th17 cells in the gut tissues and BM. Remarkably, both the LP-SI and LP-LI have significantly enhanced populations of Th17 cells (p<0.05) in the ovx group compared to the control group. A similar trend of Th17 cells was observed in the BM with a significantly higher percentage of Th17 cells (p<0.05) in the ovx group in comparison to the sham group. Strikingly LA treatment was able to significantly decrease Th17 cells in all the gut tissues i.e., LP-SI (p<0.05), LP-LI (p<0.05), MLN (p<0.01), and BM (p<0.05) **(Supplementary Fig. 5 & Fig. 6A)**. Importantly, the percentage of pTregs in the LP-SI and LP-LI was negatively correlated with the proportion of gut resident Th17 cells (**Fig. 6B)**, indicating that under post-menopausal osteoporotic conditions, pTregs could convert into Th17 cells and LA in a butyrate-mediated manner suppress this conversion.

**Figure 6.**
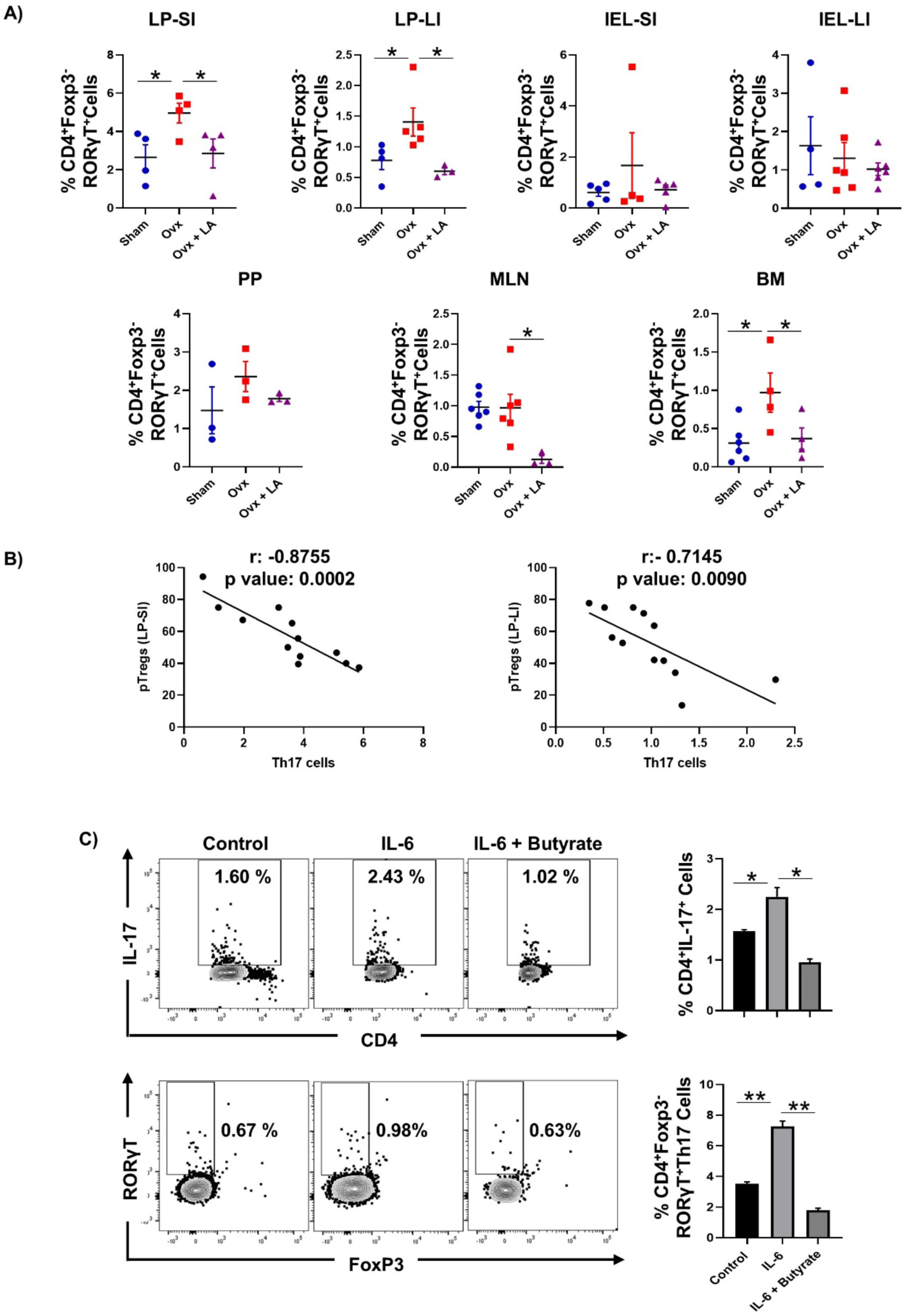
LA administration decreases gut resident Th17 cells. Cells from lamina propria of small intestine (LP-SI) and large intestine (LP-LI), intestinal epithelial lymphocytes from the small intestine (IEL-SI) and large intestine (IEL-LI), Peyer’s patches (PP), mesenteric lymph nodes (MLN), and BM of mice from sham, ovx and ovx + LA groups were harvested and analysed by flow cytometry for percentage of Th17 (CD4^+^FoxP3^-^RORγT^+^) cells. **A)** Bar graph representing percentages of Tregs in sham, ovx, and ovx + LA. **B)** Correlation graphs depicting the correlation between Th17 cells and pTregs from the LP-SI and LP-LI. The results were evaluated using the unpaired student t-test. Values are expressed as mean ± SEM (n=6) and similar results were obtained in two independent experiments. **C)** Analysis of CD4^+^IL-17^+^ cells and Th17 cells in the *in vitro* induced Tregs in the presence or absence of IL-6 and butyrate by flow cytometry. Statistical significance was defined as *p ≤ 0.05, **p < 0.01 ***p ≤ 0.001 concerning the indicated mice group.

To further validate that increased levels of IL-6 (as reported in earlier studies ^25^) results in the transdifferentiation of Tregs into Th17 cells, we isolated the naïve T cells from the MLN and cultured them in the Tregs polarizing conditions in the presence or absence of IL-6 and butyrate. We observed that the percentage of CD4^+^IL-17^+^ cells increased in the presence of IL-6. However, the addition of butyrate to the IL-6 treated cells significantly decreased the levels of CD4^+^IL-17^+^ cells **(Fig. 6C).** Similarly, the proportion of Th17 cells increased in the IL-6 treated cells. On the other hand, butyrate significantly decreased the differentiation of Th17 cells. These results further validate that increased levels of IL-6 are associated with the transdifferentiation of pTregs into exFox3 Tregs i.e. Th17 cells **(Fig. 6C).**

### 3.5 LA administration induces differentiation of RORγT^-^FoxP3^+^pTregs in ovx mice

pTregs are divided into two subsets: RORγT^-^ and RORγT^+^ pTregs. Thus, to evaluate the effect of ovariectomy on the differentiation of these two subsets, we gated pTregs for RORγT^-^ or RORγT^+^ cells **(Supplementary Fig. 1 & 2)**. To our surprise, we observed that all the tissues: gut tissues (LP-SI, LP-LI, IEL-LI, PP, and MLN), and BM have a significantly lower population of RORγT^-^ pTregs (p<0.05), but enhanced population of RORγT^+^ pTregs (p<0.05) in the ovx mice in comparison to the sham **(Supplementary Fig. 6 & Fig. 7A-B)**, highlighting the critical role of pTreg subsets in the pathophysiology of PMO. LA administration further significantly enhanced the proportion of RORγT^-^ pTregs (p<0.05) and decreased that of RORγT^+^ pTregs (p<0.05) as compared to the ovx mice in all the gut tissues (LP-SI, LP-LI, IEL-LI, PP, and MLN), and BM **(Fig. 7A-B)**. Additionally, the proportion of RORγT^-^ pTregs in the LP-SI and LP-LI was positively correlated with the BMDs of femur trabecular bone, while the frequencies of RORγT^+^ pTregs showed a negative correlation **(Fig. 7C).**

**Figure 7.**
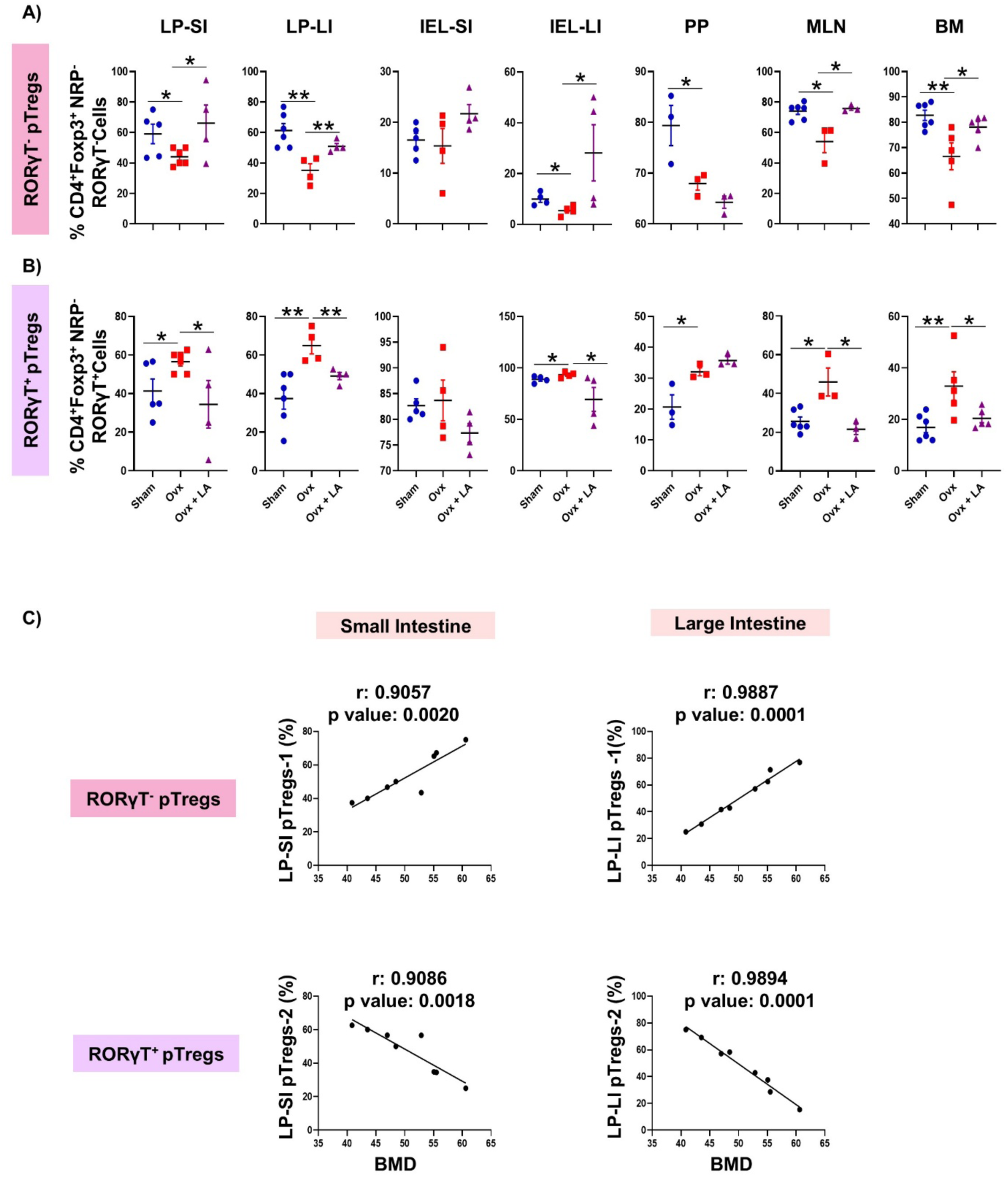
LA administration induces differentiation of RORγT^-^ pTregs *in vivo*. Cells from lamina propria of the small intestine (LP-SI) and large intestine (LP-LI), intestinal epithelial lymphocytes from the small intestine (IEL-SI) and large intestine (IEL-LI), Peyer’s patches (PP), mesenteric lymph nodes (MLN), and BM of mice from sham, ovx and ovx + LA groups were harvested and analyzed by flow cytometry for the percentage of RORγT^-^ pTregs (CD4^+^NRP^-^RORγT^-^Foxp3^+^ cells) and RORγT^+^ pTregs (CD4^+^NRP^-^RORγT^+^Foxp3^+^ cells). **A)** Bar graph representing percentages of RORγT^-^ pTregs in sham, ovx, and ovx + LA. **B)** Bar graph representing percentages of RORγT^+^ pTregs in sham, ovx, and ovx + LA. **C)** Correlation graphs depicting the correlation of RORγT^-^ pTregs and RORγT^+^ pTregs from the LP-SI and LP-LI with the bone mineral density (BMD) of femur trabecular bone. The results were evaluated using the unpaired student t-test. Values are expressed as mean ± SEM (n=6) and similar results were obtained in two independent experiments. Statistical significance was defined as *p ≤ 0.05, **p < 0.01 ***p ≤ 0.001 concerning the indicated mice group.

Since pTregs can differentiate into Th17 cells under PMO conditions, we next elucidated which pTregs subset is primarily being converted into Th17 cells. It has been demonstrated that retinoic acid (RA) can induce the expression of RORγT in Tregs and therefore, we next cultured naïve T cells under Tregs polarizing conditions in the +/-of RA (1 nM) and butyrate for 5 days ^25^ **(Fig. 8A)**. RA treatment significantly enhanced the RORγT expression in the *in vitro* cultured cells compared to the control **(Fig. 8B)**. However, butyrate treatment was not able to enhance the RORγT expression further. It is reported that RORγT stabilizes the expression of Foxp3 in Tregs. Therefore, we next analyzed the expression of Foxp3 in CD4^+^RORγT^+^ cells and observed that Foxp3 levels were significantly enhanced in the RA-treated cells, and butyrate treatment further increased the same **(Fig. 8B)**. Altogether our findings highlight that due to the absence of RORγT, RORγT^-^FoxP3^+^pTregs (in the intestine and bone marrow) show lower expression of Foxp3 and thus are more likely to transdifferentiate into ex-Foxp3 Tregs (i.e. osteoclastogenic Th17 cells).

**Figure 8.**
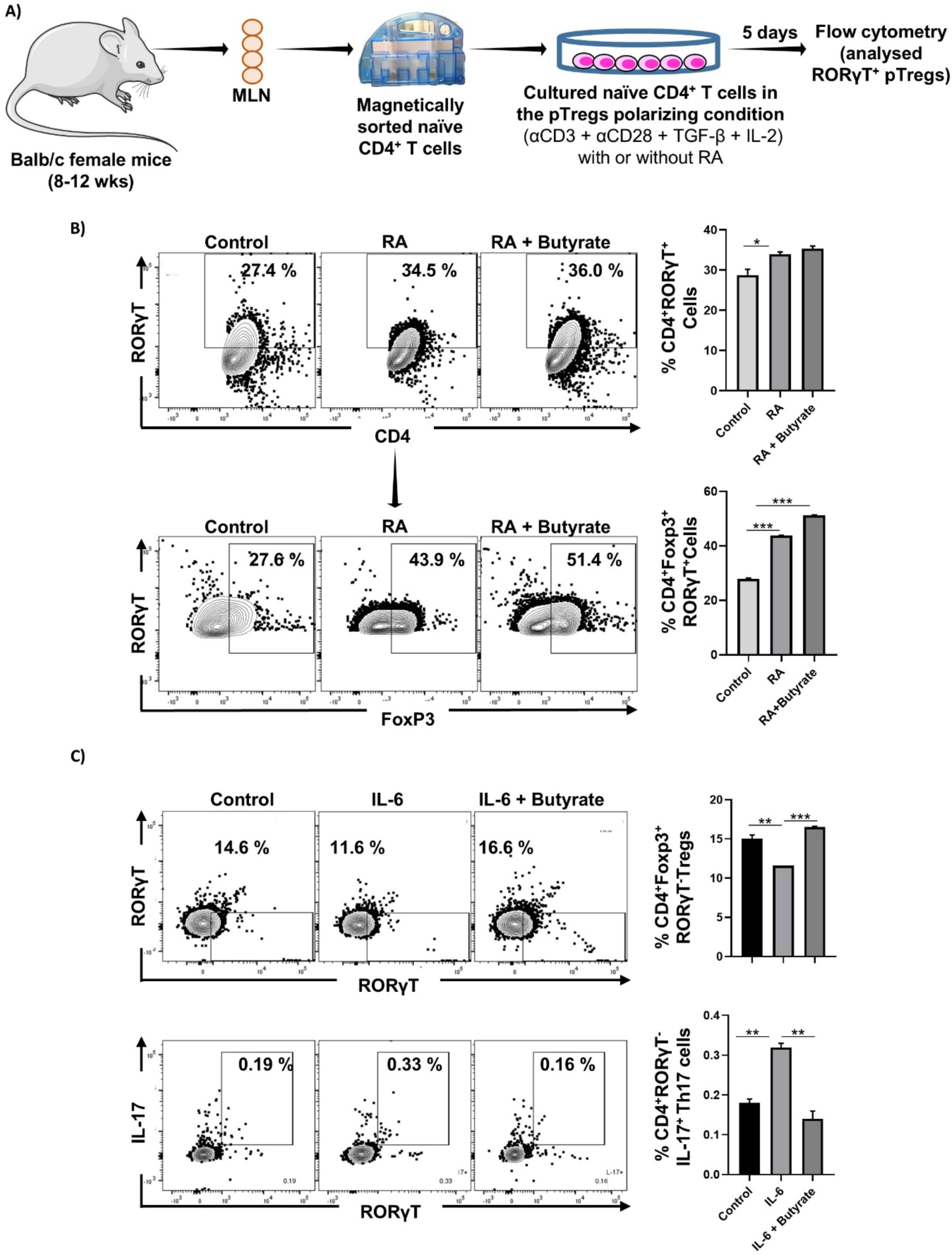
Generation of RORγT^+^ pTregs *in vitro*. **A)** Naïve T cells were isolated from the mesenteric lymph nodes (MLN) of mice and were cultured in the Tregs polarizing conditions in the presence or absence of retinoic acid with or without RA. **B)** CD4^+^RORγT^+^ cells were analyzed in the total induced cells. CD4^+^RORγT^+^ cells were further gated for Foxp3^+^ cells. Similar results were obtained in at least two independent experiments (n≥2). The results were evaluated using the unpaired student t-test. **C)** Analysis of RORγT^-^ pTregs and RORγT^+^IL-17^+^ cells in the *in vitro* induced cells in the presence of IL-6 and butyrate. Similar results were obtained in two independent experiments. Statistical significance was defined as *p ≤ 0.05, **p < 0.01 ***p ≤ 0.001 concerning the indicated mice group.

To further validate that increased levels of IL-6 in ovx mice drive the conversion of RORγT^-^ pTregs into Th17 cells we cultured the naïve T cells in Tregs polarizing conditions in the +/- of IL-6 and butyrate. We observed that RORγT^-^ pTregs were significantly decreased in the presence of IL-6 **(Fig. 8C).** However, butyrate treatment to the IL-6 treated cells significantly enhanced the differentiation of RORγT^-^ pTregs **(Fig. 8C).** We further observed that IL-6 treatment increased the proportion of CD4^+^RORγT^+^IL-17^+^ cells (i.e. Th17 cells) in the cultured cells and butyrate treatment, on the other hand, significantly decreased the proportion of these cells **(Fig. 8C).** Therefore, our results highlighted that IL-6 promotes the transdifferentiation of RORγT^-^ pTregs to Th17 cells and butyrate prevents this conversion.

### 3.6 Butyrate-primed pTregs are more potent in inhibiting osteoclastogenesis

Since butyrate enhanced pTregs, we thus next evaluated the effect of butyrate-primed pTregs on regulating the development of the osteoclasts and osteoblasts. For osteoclastogenesis, butyrate-primed pTregs (B-pTregs) were cocultured with the BMCs (BMCs: pTregs; 1:10, 1:5, and 1:1) in the presence of the MCSF and RANKL **(Fig. 9A).** On day 4, cells were stained for TRAP. The results indicated that B-pTregs were more potent in inhibiting osteoclastogenesis than the control pTregs as indicated by the significant decrease in the number of the cells with more than 3 nuclei and the area of the osteoclasts **(Fig. 9B).** For osteoblastogenesis, BMCs were cocultured with B-pTregs (BMCs: pTregs; 1:10, 1:5, and 1:1) in the presence of osteogenic induction media. On day 7, cells were processed for ALP staining. Notably, B-pTregs were unable to further enhance osteoblastogenesis compared to pTregs **(Supplementary Fig. 7)**. Altogether, butyrate modulates bone remodeling by enhancing the anti-osteoclastogenic potential of pTregs.

**Figure 9.**
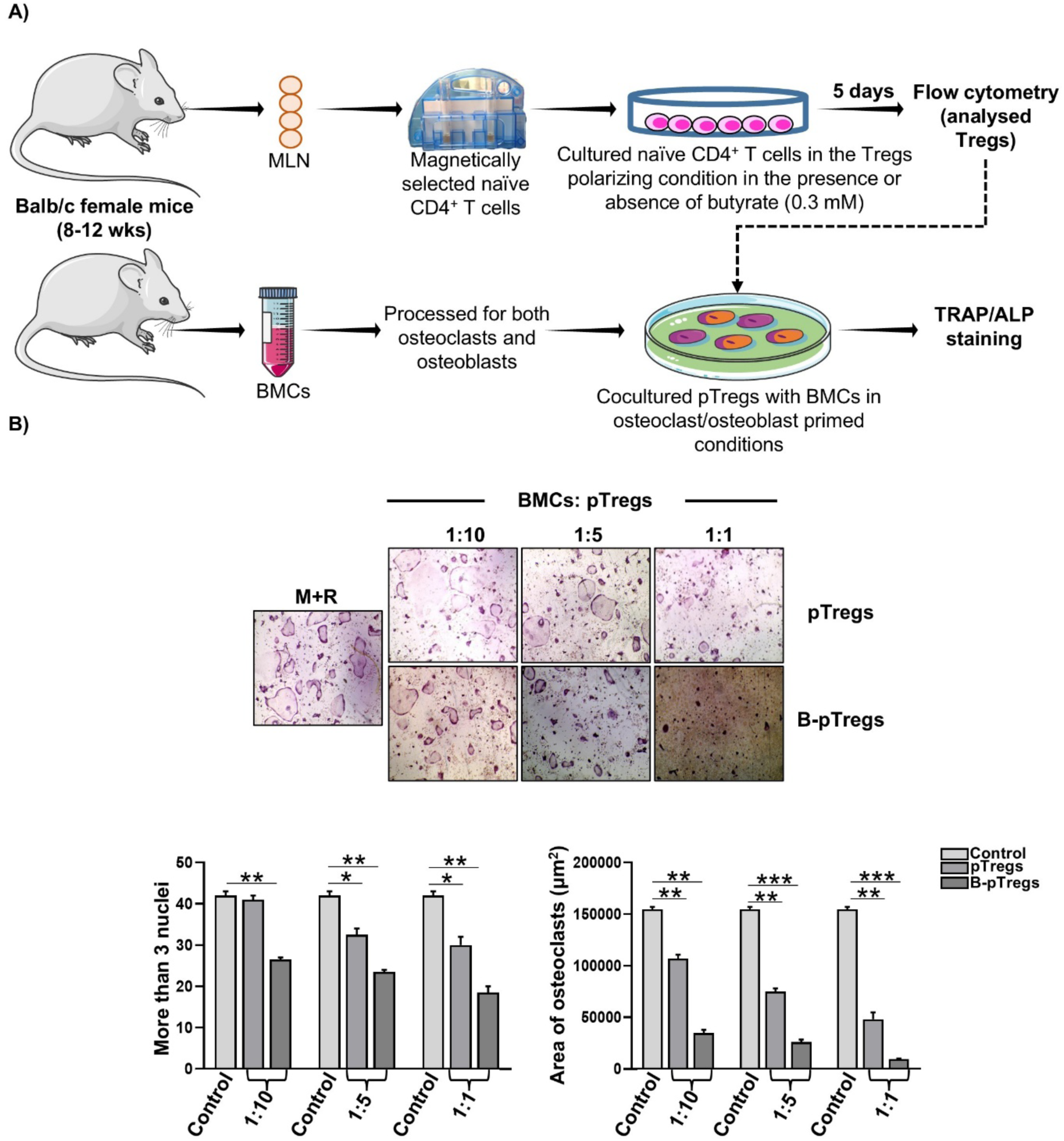
Butyrate-primed pTregs suppress osteoclastogenesis in a dose-dependent manner. A) Naïve T cells were magnetically selected from the mesenteric lymph nodes (MLN) and cultured in the Tregs polarizing conditions in the presence and absence of 0.3 mM butyrate After incubation of 5 days pTregs were cocultured with the bone marrow cells (BMCs) in the presence of M-CSF (30ng/ml) and RANKL (60 ng/ml) for 4 days. B). Osteoclast differentiation was induced in BMCs with M-CSF (30/ml) and RANKL (60 ng/ml) with pTregs or butyrate-primed pTregs at different ratios for 4 days. Giant multinucleated cells were stained with TRAP, and cells with ≥ 3 nuclei were considered as mature osteoclasts. Photomicrographs at 10X magnification were taken. Osteoclast differentiation was analyzed by plotting for the number of TRAP-positive cells with more than 3 nuclei and an area of osteoclasts. Similar results were obtained in at least two independent experiments (n≥2). Statistical significance was considered as p≤0.05(*p≤0.05, **p≤0.01, ***p≤0.001) with respect to indicated groups.

## 4.0 Discussion

Osteoporosis is a prevalent inflammatory bone loss disorder that affects over 500 million people globally ^5^. Our lab has proposed the term “Immunoporosis”, specifically referring to the pivotal role of immune cells in the pathophysiology of osteoporosis ^5–8^. The function of Tregs in maintaining bone homeostasis has been thoroughly explained in this respect ^9–13^. Tregs are bone-protective cells and secrete several anti-osteoclastogenic cytokines, such as TGF-β and IL-10 ^14–16^. The gut is the largest reservoir of the Tregs ^18^. Studies in the past decade have highlighted the importance of a healthy gut in bone homeostasis, as dysbiosis in the gut and loss of intestinal permeability are directly associated with bone loss ^26,27^. However, no study has shown the role of GTregs in regulating bone health. Therefore, we studied the effect of LA in modulating the GTregs. Interestingly, our flow cytometry data demonstrated that ovx mice have a significantly decreased population of GTregs, highlighting their crucial role in regulating bone health, and administration of LA reversed this trend. GTregs are further subdivided into two subsets, viz., pTregs and tTregs, and several studies have explored the role of these two subsets of Tregs in various pathologies ^28^. However, the role of pTregs and tTregs in modulating bone health is still unexplored. Thus, our group, for the first time, delved into exploring the role of these two Treg subpopulations in PMO. Fascinatingly, we observed that the composition of pTregs and tTregs is significantly altered in PMO and that LA restores the same via inducing the differentiation of pTregs. Several previous studies have pointed out that gut-associated metabolites (viz. SCFAs) have a significant role in the induction of pTregs, and *Lactobacillus* species have established roles in enhancing the production of SCFAs ^21^. Therefore, we next performed metabolomics (fecal samples) to analyze the status of SCFAs in our preclinical model of PMO. Our data demonstrated that ovx mice have significantly lower levels of SCFAs (acetate, propionate, and butyrate) and that LA administration significantly enhanced the levels of butyrate in the ovx mice. Interestingly, our results further indicated that butyrate significantly enhanced the differentiation of GTregs (explicitly that of the pTregs rather than the tTregs).

Further, we were inquisitive to find the mechanism involved in butyrate-mediated enhancement of pTregs. Earlier studies reported that the stability of Foxp3 expression affects the balance of tolerance and autoimmunity, as well as the efficacy of Treg cell-based therapy^24^. The fate of plastic Foxp3^+^ Treg cells is modulated by various inflammatory conditions, such as IL-6 and TNFα, which further induce transdifferentiation of Tregs into Th17 cells ^22–24^. Fate mapping studies indicated that pTregs are more prone to lose Foxp3 expression as tTregs have a more stable phenotype ^22,29^. Tregs that lose FoxP3 expression are called ex-Foxp3 Tregs (i.e. Th17 cells recognized by higher expression of IFN-γ and IL-17) ^22^. Since pTregs were significantly decreased in ovariectomized mice we are thus interested in determining the status of transdifferentiation of pTregs into exFoxp3 Tregs under PMO conditions. For the same serum cytokine analysis was performed which revealed higher levels of inflammatory cytokines viz. IFN-γ, TNFα, and IL-17, and decreased levels of IL-10 in the ovariectomized mice, thereby promoting the transdifferentiation of pTregs to exFoxp3 Tregs. Remarkably, LA administration significantly reversed this cytokine imbalance. Our *in vitro,* analysis further indicated that butyrate diminished the production of inflammatory cytokines IFN-γ and IL-17 from pTregs without affecting that from tTregs. Moreover, butyrate averts the generation of Th17 cells from Tregs under both presence or absence of IL-6, further corroborating our *in vivo* data where in Th17 cells are enhanced in ovx mice and LA administration decreased the same. The frequency of pTregs in gut tissues was observed to be negatively correlated with Th17 cells. pTregs are further divided into two subsets viz. RORγT^-^ and RORγT^+^ pTregs, RORγT^-^ pTregs are induced by food antigens and represent a subpopulation that can be distinguished from RORγT^+^ pTregs (induced in response to gut microbiota). We thus next determined which pTreg subset is more inclined to transdifferentiate towards Th17 cells. We observed that it’s the RORγT^-^ pTregs that were significantly decreased in the ovx mice, and LA treatment enhanced their population, thereby highlighting the possibility of RORγT^-^ pTregs being converted to Th17 cells under PMO conditions. To validate this, we generated pTregs *in vitro* in the presence of RA (known to induce RORγT^+^ pTregs) ^25^. We further observed that RA-primed Tregs showed higher expression of FoxP3 compared to the unprimed Tregs and thus represent a more stable phenotype. Moreover, our results indicate that IL-6 promotes the transdifferentiation of pTregs into Th17 cells and butyrate prevents this conversion. Altogether, our study for the first time demonstrated that Th17 cells under PMO conditions develop from RORγT^-^ pTregs. Also, our findings reveal that the “pTreg-Th17” cell axis does play a pivotal role in the pathophysiology of PMO, and probiotic-LA mitigates PMO-induced bone loss via augmenting the immunoporotic potential of pTregs.

## Acknowledgment

AB, LS, and RKS acknowledge the Department of Biotechnology, AIIMS, New Delhi-India for providing infrastructural facilities. AB and LS thank ICMR for the research fellowship. A graphical abstract is created with the help of Servier Medical Art, provided by Servier, licensed under a Creative Commons Attribution 3.0 unported license (https://smart.servier.com).

## Credit authorship contribution statement

RKS contributed to the conception and design of the study; AB and LS contributed to the acquisition and analysis of data; AB contributed to drafting the text or preparing the figures. AT helped with SEM analysis. PKM provided valuable inputs. All authors reviewed the manuscript. All authors contributed to the article and approved the submitted version.

