## Supplementary Figures for "*Lactobacillus acidophilus* ameliorates inflammatory bone loss under post-menopausal osteoporotic conditions via preventing the pathogenic conversion of gut resident pTregs into Th17 cells"

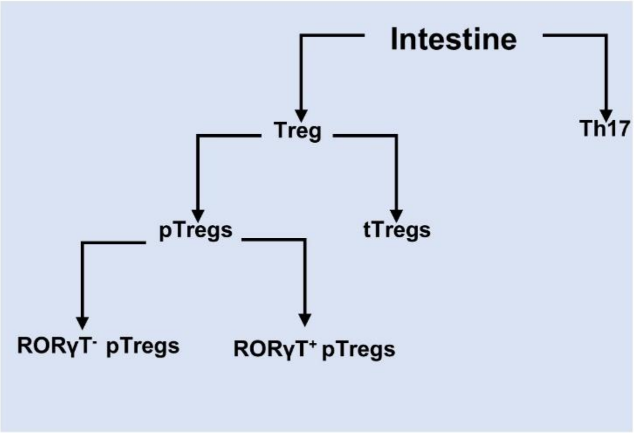


| Immune cell | Phenotype |
| --- | --- |
| **Tregs** | CD4$^+$FoxP3$^+$ |
| **tTregs** | CD4$^+$FoxP3$^+$NRP$^+$ |
| **pTregs** | CD4$^+$FoxP3$^+$NRP$^-$ |
| **RORγT$^-$ pTregs** | CD4$^+$FoxP3$^+$NRP$^-$RORγT$^+$ |
| **RORγT$^+$ pTregs** | CD4$^+$FoxP3$^+$NRP$^-$RORγT$^-$ |
| **Th17 cells** | CD4$^+$Foxp3$^-$RORγT$^+$ |

**Supplementary Figure 1.** Flow chart representing the immune cells harvested from the lamina propria of small (LP-SI) and large intestine (LP-LI) and phenotypes used for analysis of these immune cells by flow cytometry

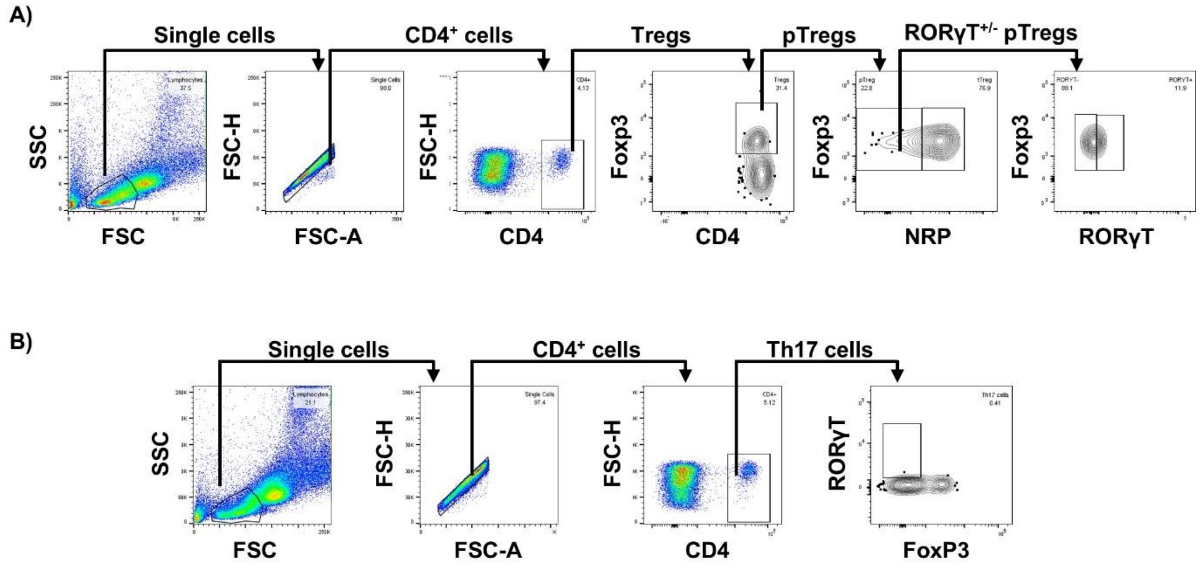


**Supplementary Figure 2.** Gating strategy used for the analysis of (A) Tregs and (B) Th17 cells.

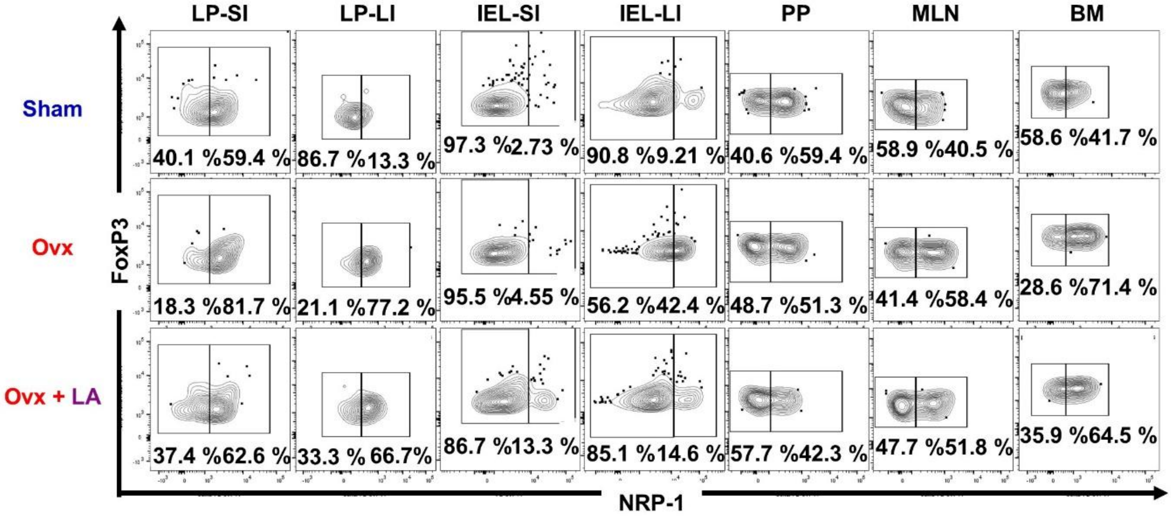


**Supplementary Figure 3. LA administration modulates gut resident pTregs and tTregs *in vivo*).** Cells from lamina propria of the small intestine (LP-SI) and large intestine (LP-LI), intestinal epithelial lymphocytes from the small intestine (IEL-SI) and large intestine (IEL-LI), Peyer's patches (PP), mesenteric lymph nodes (MLN), and BM of mice from sham, ovx and ovx + LA groups were harvested and analyzed by flow cytometry for the percentage of pTregs ($CD4^+NRP^-Foxp3^+$ cells) and tTregs ($CD4^+NRP^+Foxp3^+$ cells

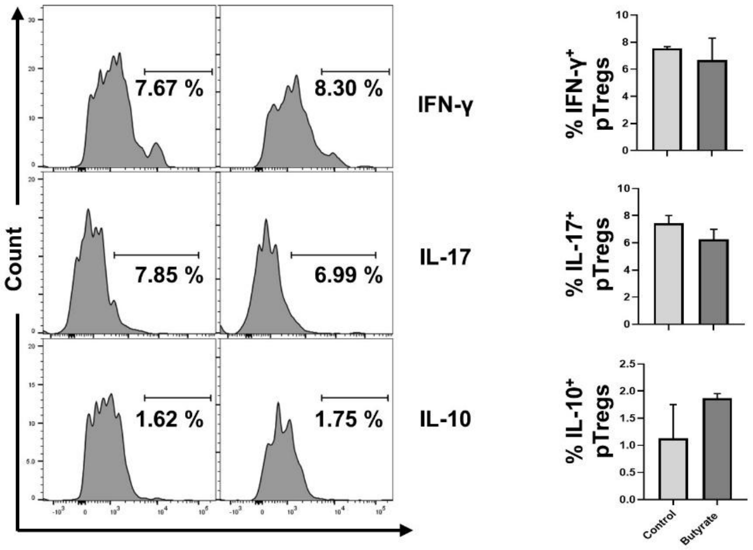


**Supplementary Figure 4. Butyrate does not modulate the cytokine secretion from tTregs.** Analysis of IFN-γ, IL-17, and IL-10 levels in the tTregs population by flow cytometry in the presence and absence of butyrate. Statistical significance was considered as $p \leq 0.05$($*p \leq 0.05$,$**p \leq 0.01$, $***p \leq 0.001$) with respect to indicated groups.

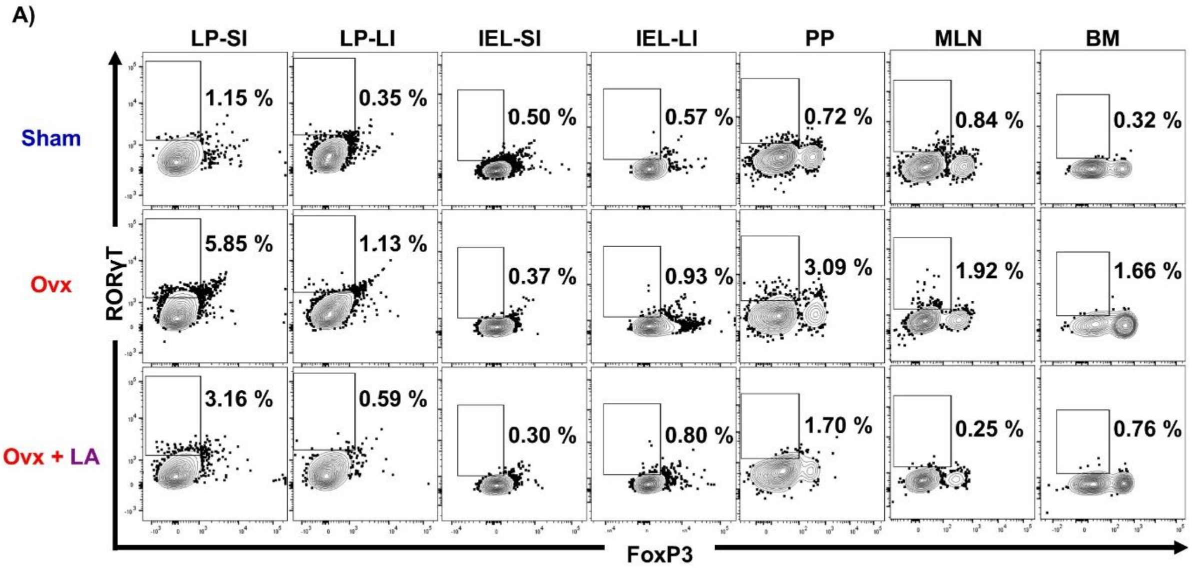


**Supplementary Figure 5. LA administration decreases Th17 cells *in vivo*.** Cells from lamina propria of small intestine (LP-SI) and large intestine (LP-LI), intestinal epithelial lymphocytes from the small intestine (IEL-SI) and large intestine (IEL-LI), Peyer's patches (PP), mesenteric lymph nodes (MLN), and BM of mice from sham, ovx and ovx + LA groups were harvested and analysed by flow cytometry for percentage of Th17 ($CD4^{+}FoxP3^{-}ROR\gamma T^{+}$) cells.

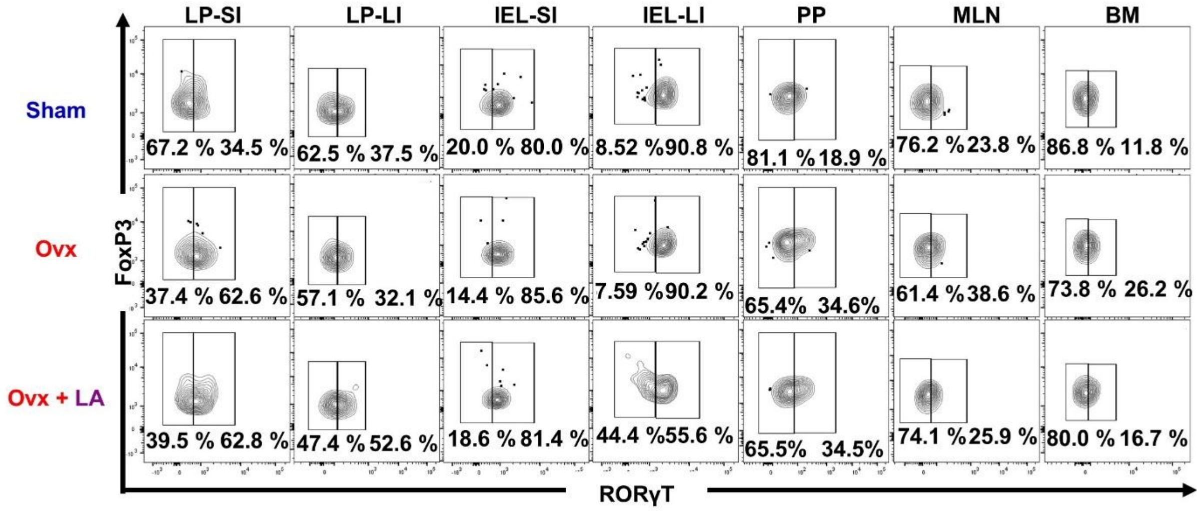


**Supplementary Figure 6. LA administration induces differentiation of RORγT⁻ pTregs *in vivo*.** Cells from lamina propria of the small intestine (LP-SI) and large intestine (LP-LI), intestinal epithelial lymphocytes from the small intestine (IEL-SI) and large intestine (IEL-LI), Peyer's patches (PP), mesenteric lymph nodes (MLN), and BM of mice from sham, ovx and ovx + LA groups were harvested and analyzed by flow cytometry for the percentage of RORγT$^-$ pTregs (CD4$^+$NRP$^-$RORγT$^-$Foxp3$^+$ cells) and RORγT$^{-+}$pTregs (CD4$^+$NRP$^-$RORγT$^+$Foxp3$^+$ cells).

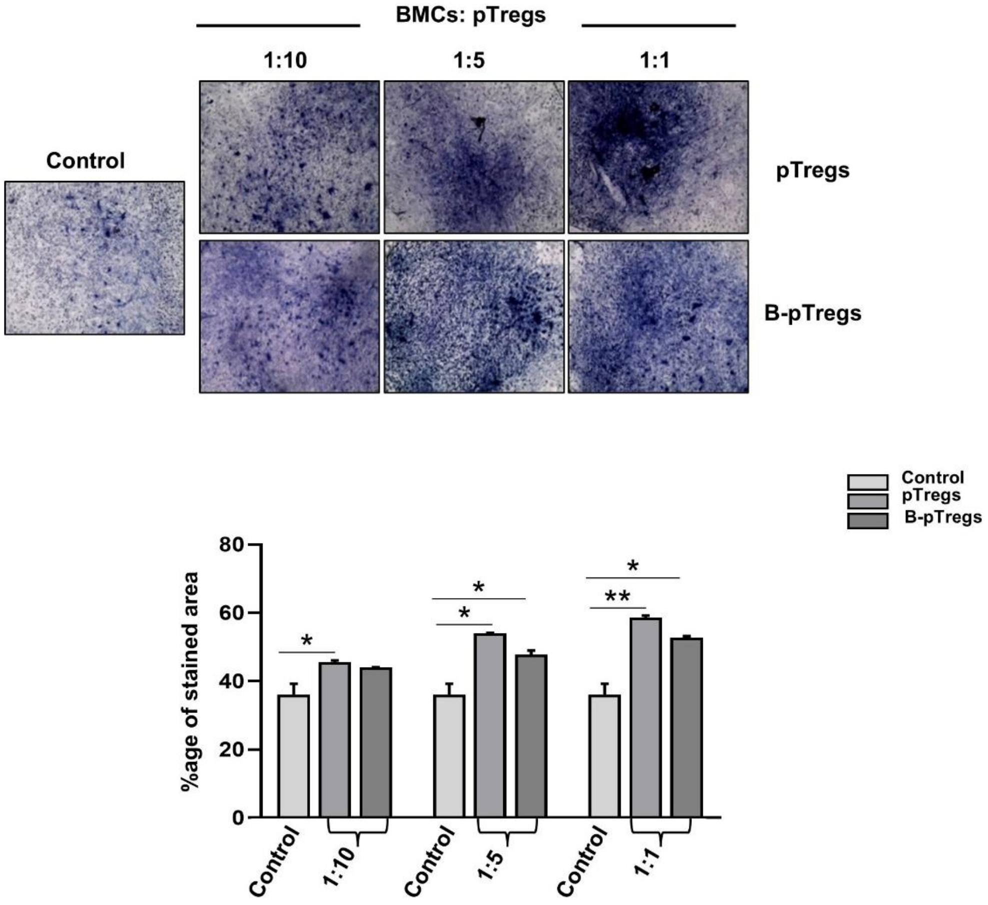


**Supplementary Figure 7. Butyrate-primed pTregs do not affect osteoblastogenesis.** Naïve T cells were magnetically selected from the mesenteric lymph nodes (MLN) and cultured in the Tregs polarizing conditions in the presence and absence of 0.3 mM butyrate After incubation of 5 days pTregs were cocultured with the bone marrow cells (BMCs) in the presence of osteogenic induction media. Osteoblastogenesis was determined by alkaline phosphatase (ALP) staining and the percentage of the stained area was calculated with the help of ImageJ software. Similar results were obtained in at least two independent experiments ($n \geq 2$). Statistical significance was considered as $p \leq 0.05$(*$p \leq 0.05$,**$p \leq 0.01$, ***$p \leq 0.001$)with respect to indicated groups.
